# Destabilization of chromosome structure by histone H3 lysine 27 methylation

**DOI:** 10.1101/454223

**Authors:** Mareike Möller, Klaas Schotanus, Jessica Soyer, Janine Haueisen, Kathrin Happ, Maja Stralucke, Petra Happel, Kristina M. Smith, Lanelle R. Connolly, Michael Freitag, Eva H. Stukenbrock

## Abstract

Chromosome and genome stability are important for normal cell function as instability often correlates with disease and dysfunction of DNA repair mechanisms. Many organisms maintain supernumerary or accessory chromosomes that deviate from standard chromosomes. The pathogenic fungus *Zymoseptoria tritici* has as many as eight accessory chromosomes, which are highly unstable during meiosis and mitosis, transcriptionally repressed, show enrichment of repetitive elements, and enrichment with heterochromatic histone methylation marks, e.g., trimethylation of H3 lysine 9 or lysine 27 (H3K9me3, H3K27me3). To elucidate the role of heterochromatin on genome stability in *Z. tritici,* we deleted the genes encoding the methyltransferases responsible for H3K9me3 and H3K27me3, *kmt1* and *kmt6*, respectively, and generated a double mutant. We combined experimental evolution and genomic analyses to determine the impact of these deletions on chromosome and genome stability, both *in vitro* and *in planta*. We used whole genome sequencing, ChIP-seq, and RNA-seq to compare changes in genome and chromatin structure, and differences in gene expression between mutant and wildtype strains. Analyses of genome and ChIP-seq data in H3K9me3-deficient strains revealed dramatic chromatin reorganization, where H3K27me3 is mostly relocalized into regions that are enriched with H3K9me3 in wild type. Many genome rearrangements and formation of new chromosomes were found in the absence of H3K9me3, accompanied by activation of transposable elements. In stark contrast, loss of H3K27me3 actually increased the stability of accessory chromosomes under normal growth conditions *in vitro*, even without large scale changes in gene activity. We conclude that H3K9me3 is important for the maintenance of genome stability because it disallows H3K27me3 in these regions. In this system, H3K27me3 reduces the overall stability of accessory chromosomes, generating a “metastable” state for these quasi-essential regions of the genome.

**Author Summary:** Genome and chromosome stability are essential to maintain normal cell function and viability. However, differences in genome and chromosome structure are frequently found in organisms that undergo rapid adaptation to changing environmental conditions, and in humans are often found in cancer cells. We study genome instability in a fungal pathogen that exhibits a high degree of genetic diversity. Regions that show extraordinary diversity in this pathogen are the transposon-rich accessory chromosomes, which contain few genes that are of unknown benefit to the organism but maintained in the population and thus considered “quasi essential”. Accessory chromosomes in all fungi studied so far are enriched with markers for heterochromatin, namely trimethylation of H3 lysine 9 and 27 (H3K9me3, H3K27me3). We show that loss of these heterochromatin marks has strong but opposing effects on genome stability. While loss of the transposon-associated mark H3K9me3 destabilizes the entire genome, presence of H3K27me3 favors instability of accessory chromosomes. Our study provides insight into the relationship between chromatin and genome stability and why some regions are more susceptible to genetic diversity than others.

## Introduction

Chromatin structure plays an important role in genome organization and gene expression [1–3]. A well-studied hallmark of epigenetic regulation is the reversible modification of histone tails, which can alter chromatin structure [4]. Chromatin structure determines accessibility of the underlying DNA to regulatory elements, whereby tightly packed DNA, known as heterochromatin, is less accessible for DNA binding proteins and usually shows little transcriptional activity [5]. Heterochromatic regions often cluster together and are spatially separated from more transcriptionally active and accessible euchromatic regions [6]. Specific histone modifications are associated with either heterochromatic or euchromatic regions. Some of the most studied histone modifications are histone H3 lysine 9 di‐ or trimethylation (H3K9me2/3) and H3K27me2/3 as markers for heterochromatin and H3K4me2/3 as markers for euchromatin [7].

H3K9me2/3 is catalyzed by the histone methyltransferase KMT1 (Su[var]3-9) [8,9], in fungi also called Clr4 [10] or DIM-5 [11]. Previous studies demonstrated enrichment of this constitutive heterochromatin mark in repeat-rich regions and a clear link with the control of transposable elements (TE) and genome stability [12–14]. H3K9me2/3 have been shown to be involved in suppression of meiotic recombination in *Arabidopsis thaliana* [15] and the control of DNA methylation in *Neurospora crassa* [11].

H3K27me2/3, associated with “facultative heterochromatin”, is catalyzed by KMT6 (E[Z]) as part of the PRC2 complex [16]. In plants, fungi, and animals, this histone mark is used to generate “transcriptional memory” and is easily reversible when environmental or endogenous stimuli require organismal responses. In many organisms, H3K27 methylation is required for development and cell differentiation [17–23], and aberrant H3K7me3 distribution is prevalent in cancer cells [24–26]. In fungi, H3K27me3 correlates with subtelomeric gene silencing [22,23,27], and has been shown to play a role in development, pathogenicity, and transcriptional regulation of secondary metabolite gene clusters [21,28,29].

H3K27me3 is also a hallmark of accessory chromosomes, which are found in several fungal plant pathogens [28,30,31]. Accessory chromosomes are not essential for survival under all environmental conditions, and thus encode “quasi-essential” genes [32] that can confer selective advantages under some conditions e.g. in a specific host species, resulting in presence or absence of these chromosomes among specific individuals of a given species. They are also characterized by extensive structural rearrangements and length variation [33,34]. In some species (*Fusarium oxysporum*, *Nectria haematococca*, *Alternaria alternata*), accessory chromosomes increase virulence [35–38]. However, in the wheat pathogen *Zymoseptoria tritici*, some accessory chromosomes have been demonstrated to confer reduced fitness and virulence *in planta* [39], suggesting that there are other stages in the life cycle when they become important. Accessory chromosomes of fungi differ structurally from core chromosomes by higher repeat and lower gene density compared to core chromosomes and show little transcriptional activity [35,40–43]. Transcriptional silencing can be explained by their predominantly heterochromatic structure, with H3K27me3 enrichment on almost the entire chromosome and H3K9me3 covering repetitive sequences [28,30]. Centromeres and telomeres are important structural components of chromosomes. In plants, centromeres of B chromosomes, equivalents to fungal accessory chromosomes, differ from those of A chromosomes [44], but in *Z. tritici* centromeres, telomere repeats, and subtelomeric regions are so far by all measures near identical on core and accessory chromosomes [31]. Though accessory chromosomes are a frequent phenomenon in fungi, little is known about their origin and maintenance. Studies on chromosome stability revealed that accessory chromosomes are highly unstable, both during mitosis [36,45,46] and meiosis [47].

Here we investigated to what extent the particular histone methylation pattern on accessory chromosomes contributes to the structural differences, transcriptional repression and instability. We shed light on the roles of H3K9me3 and H3K27me3 on genome stability in the hemi-biotrophic wheat pathogen *Z. tritici* that reproduces both asexually and sexually. By combining experimental evolution with genome, transcriptome and ChIP sequencing, we show that both heterochromatin-associated histone methylation marks contribute significantly, but in distinct ways, to chromosome stability and integrity. While the presence of H3K27me3 enhances chromosome loss and instability, loss of H3K9me3 promotes chromosome breakage, segmental duplications as well as the formation of new chromosomes − possibly resembling the emergence of accessory chromosomes. Taken together, our findings demonstrate the importance of constitutive heterochromatin for maintaining genome stability and gene silencing as well as an unexpected destabilizing influence of facultative heterochromatin on mitotic accessory chromosome transmission. The presence of eight accessory chromosomes in the reference isolate IPO323 makes *Z. tritici* an excellent model to study accessory chromosome characteristics and dynamics, which relates to general interest in chromosome maintenance in cancer or other aneuploid cell types.

## Results

### Deletion of histone methyltransferase encoding genes *kmt1* and *kmt6* in *Zymoseptoria tritici*

To investigate the impact of heterochromatin on fitness, transcription and genome stability in *Z. tritici*, we generated mutants of two histone methyltransferases Kmt1 (*S. pombe* Clr4; *N. crassa* DIM-5, *Fusarium* KMT1, *H. sapiens* SUV39H1) and Kmt6 (*N. crassa* SET-7; *Fusarium* KMT6; *H. sapiens* EZH2). We identified the *Z. tritici* genes by BLAST searches with the *N. crassa* and *F. graminearum* protein coding sequences as baits. Kmt1 is encoded by *kmt1* (Zt_chr_1_01919), and Kmt6 is encoded by *kmt6* (Zt_chr_4_00551) [48]. We used *Agrobacterium tumefaciens*-mediated transformation [49] to delete both genes in a derivate of the *Z. tritici* reference isolate IPO323 that lost chromosome 18 during *in vitro* growth, here called Zt09 [31,40,41]. Correct integration of the *hph* gene, which confers hygromycin resistance [49], and *kmt1* or *kmt6* deletion were verified by PCR and Southern analyses (Fig S1). We generated a double deletion mutant by deleting the *kmt1* gene in a *kmt6* deletion mutant background by using resistance to nourseothricin conferred by the *nat* gene [50] as an additional selection marker. We isolated several independent transformants, including eight *Δkmt1*, six *Δkmt6* and ten *Δkmt1 Δkmt6* double mutants (from here on abbreviated *Δk1/k6*). For further studies we selected two or three mutants of each type (Table S1). *Δkmt1* and *Δkmt6* single mutants were complemented by re-integrating the previously deleted gene and a *neo*^+^ resistance marker that can confer G418 resistance at the native gene loci (Fig S1).

We performed ChIP-seq on Zt09, *Δkmt1* (Zt125−#68, −#80), *Δkmt6* (Zt110xyh#283, −#285, −#365) and the double deletion mutant *Δk1/k6* (Zt219−#23, −#116), which verified the absence of H3K9me3 in *Δkmt1* and *Δk1/k6*, and the absence of H3K27me3 in *Δkmt6* and *Δk1/k6* mutants (Fig S2), confirming that Kmt1 and Kmt6 are the only histone methyltransferases in *Z. tritici* responsible for H3K9 and H3K27 trimethylation, respectively.

### Deletion of *kmt1*, but not *kmt6*, severely impacts *in vitro* and *in planta* growth

To assess if deletion of *kmt1* and *kmt6* has an impact on *in vitro* growth or pathogenicity on wheat, we performed comparative growth and virulence assays comparing the mutants to the wild type Zt09. To compare growth rates, the reference strain Zt09, deletion and complemented strains were grown in liquid YMS cultures and the OD_600_ was measured until cells reached stationary phase. Overall, the Δ*kmt1* strains and Δ*k1/k6* double deletion mutants showed significantly reduced growth *in vitro* (Fig S3). The Δ*kmt6* mutants and both *kmt1^+^*and *kmt6^+^*complementation strains showed no significant differences in growth compared to Zt09 (Wilcoxon rank-sum test, *p*-values: Δ*kmt1* 0.025; Δ*kmt6* 0.42; Δ*k1/k*6 0.005; *kmt1^+^*0.28; *kmt6^+^*0.63).

We furthermore assessed the tolerance of the Δ*kmt1*, Δ*kmt6* and Δ*k1/k6* mutants to abiotic stress *in vitro* by testing temperature, cell wall, oxidative, and genotoxic stressors. As observed in the growth assays, the Δ*kmt1* and Δ*k1/k6* double deletion mutants showed overall reduced growth under all tested conditions (Fig S4), especially under osmotic stress induced by high sorbitol concentrations or the cell wall-interfering agent Congo Red. The Δ*kmt6* mutants showed little differences compared to Zt09; however, elevated temperatures led to increased melanization in the Δ*kmt6* mutants suggesting involvement of H3K27me3 in the response to temperature stress. This phenotype was reversed in the complemented *kmt6^+^*strain (Fig S5).

To study the effect of the histone methyltransferase deletions on the ability to infect wheat (*Triticum aestivum*), we inoculated leaves of the susceptible cultivar Obelisk with single cell cultures of Δ*kmt1*, Δ*kmt6,* the Δ*k1/k6* double deletion mutant and Zt09. The infection assays demonstrated significant impact of both H3K27me3 and H3K9me3 on virulence. While the number of pycnidia and necrotic leaf areas only decreased in the Δ*kmt6* mutants, wheat infection by Δ*kmt1* and Δ*k1/k6* mutants resulted in almost no symptoms (Fig S6). If any symptoms developed, these appeared considerably later than symptoms caused by the reference Zt09 and the Δ*kmt6* mutants (Fig S6).

### Loss of H3K9me3 allows H3K27me3 to invade repeat-rich regions

We next addressed how the deletion of *kmt1* and *kmt6* impacts the distribution of three histone modifications (H3K4me2, H3K9me3, H3K27me3) by ChIP-seq (Table S2). We previously found that H3K4me2 is associated with gene-rich, transcriptionally active regions on core chromosomes, that constitutive heterochromatin, enriched with H3K9me3, forms almost exclusively on repetitive elements, and that facultative heterochromatin, enriched with H3K27me3, forms nearly on the entire length of all accessory chromosomes and the subtelomeric regions of core chromosomes [31]. As expected, analyses of ChIP-seq data confirmed the complete loss of H3K9me3 in the Δ*kmt1* mutant, loss of H3K27me3 in the Δ*kmt6* mutant and the absence of both marks in the Δ*k1/k6* mutants (Fig S2).

We computed the sequence coverage of each histone modification per chromosome to estimate the global effects on chromatin structure. The absence of one histone methylation mark had differential effects on the distribution of the other two methylation marks on core and accessory chromosomes (Fig 1, Table 1). In the Δ*kmt1* mutants, the amount of sequences enriched with H3K27me3 decreases on the accessory chromosomes when compared to Zt09, representing the opposite trend to the observations made on the core chromosomes, where we observed an increased amount of sequences enriched for H3K27me3 (Fig 1, Table 1). However, this effect varies on different accessory chromosomes (Table 1, Fig S7). The difference in H3K27me3 distribution can be explained by relocation of H3K27me3 to former H3K9me3-associated sequences in the *Δkmt1* mutant (Fig 1). While fewer genes are associated with H3K27me3 (Fig S7), more transposable elements (TEs) show H3K27me3 enrichment in the Δ*kmt1* mutant (Fig S7) compared to Zt09. These observations reveal that loss of H3K9me3 promotes H3K27me3 relocation to transposable elements and confers simultaneous loss of H3K27me3 at positions with this histone mark in the reference strain. The subtelomeric H3K27me3 enrichment, however, is not affected by this relocation, which explains why we observe opposite effects on core and accessory chromosomes, as core chromosomes predominantly show H3K27me3 enrichment in subtelomeric regions while accessory chromosomes show overall enrichment with H3K27me3. H3K4me2 increases on both core and accessory chromosomes, with accessory chromosomes showing a considerably higher relative increase compared to H3K4me2 in Zt09 (Table 1).

**Fig 1.**
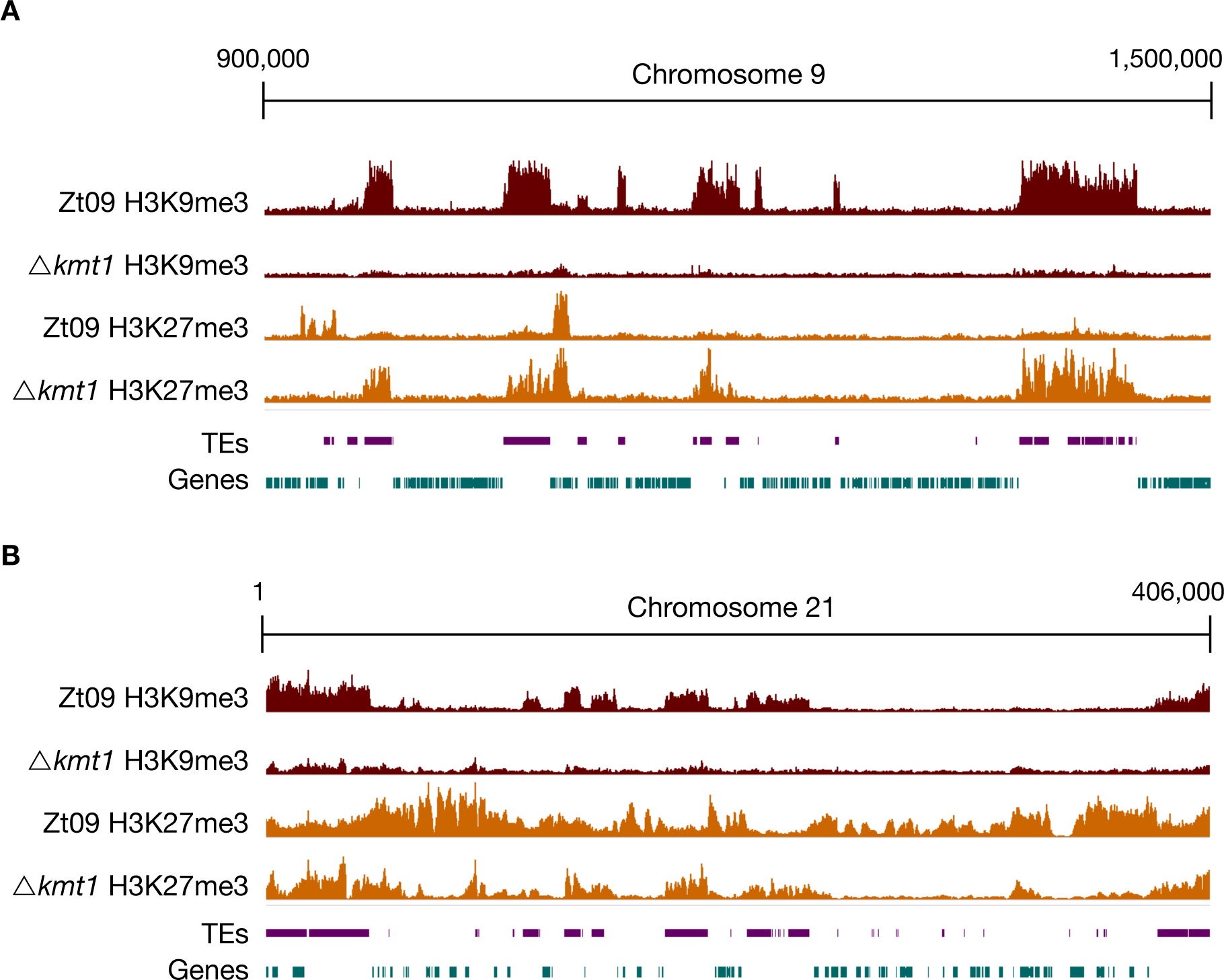
ChIP-seq reveals relocation of H3K27me3 on core (A) and accessory (B) chromosomes in Δ*kmt1* mutants. By analyzing ChIP-seq data in the Δ*kmt1* mutants we found that enrichment of H3K27me3 moves to sequences that are normally enriched with H3K9me3. A region on core chromosome 9 **(A)** is shown, where H3K27me3 is strongly enriched at former H3K9me3 regions, but depleted from its original positions. On accessory chromosomes **(B)**, here full-length chromosome 21 as an example, there are similar dynamics as observed on core chromosomes. Accessory chromosomes normally show overall enrichment of H3K27me3. In absence of H3K9me3, H3K27me3 concentrates on former H3K9me3 regions, again being depleted from its original position. However, this effect varies between accessory chromosomes (S7 Figure). The low amount of background found in Δ*kmt1* is due to the repetitive nature of the H3K9me3-enriched regions. All shown ChIP-seq tracks are normalized to RPKM [115].

**Table 1.**
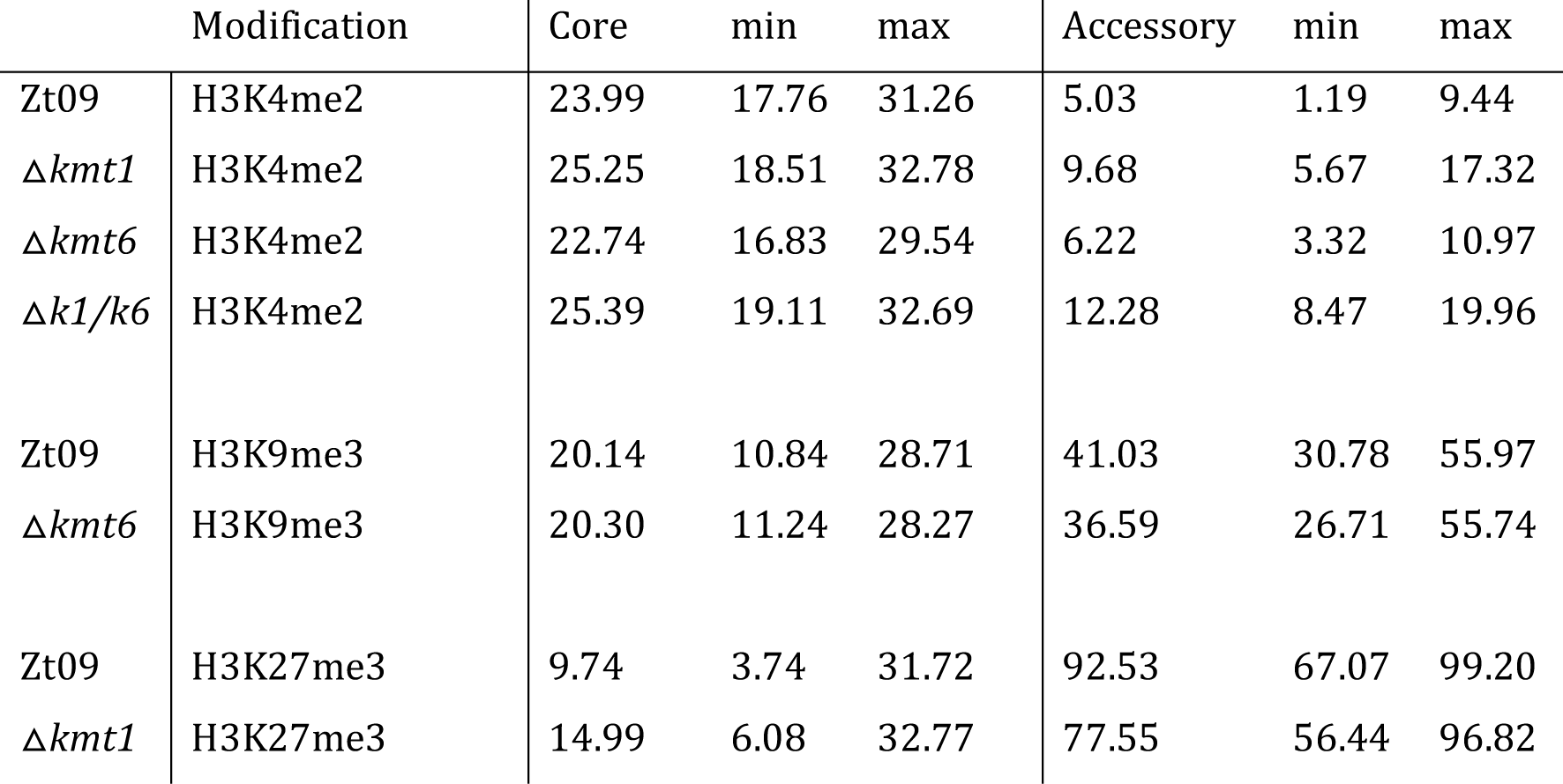
Percentage of sequence coverage (significantly enriched regions) of core and accessory chromosomes with H3K4me2, H3K9me3 and H3K27me3 relative to the chromosome length. Minimum and maximum values refer to the chromosomes showing highest or lowest sequence coverage with enrichment of the respective histone modification. H3K4me2 coverage on accessory chromosomes increases in all mutant strains, while there are little differences in the overall coverage with H3K9me3 between Zt09 and *Δkmt6*. H3K27me3 enrichment increases on core chromosomes and decreases on accessory chromosomes in the *Δkmt1* mutant.

Conversely, H3K9me3 is not affected by loss of H3K27me3 in the Δ*kmt6* mutants; there is no relocation and minor differences in coverage. H3K4me2 enrichment does increase on accessory chromosomes, but not to the same extent as observed in the Δ*kmt*1 mutants and it slightly decreases on core chromosomes (Table 1 and Fig S7), suggesting minor effects of *Δkmt6* on transcriptional activation. In the Δ*k1/k6* double deletion mutants, where both H3K9me3 and H3K27me3 are not present, we detected an increase in H3K4me2, similar to the Δ*kmt*1 single mutants on core chromosomes and slightly higher on the accessory chromosomes.

In summary, loss of H3K9me3 has a great impact on H3K27me3 distribution, while loss of H3K27me3 has little influence on H3K9me3. Deletion of *kmt1* promotes large scale relocalization of other histone modifications, indicating more dramatic effects on genome organization and transcriptional activation than deletion of *kmt6.*

### H3K27me3 has little effects on transcriptional activation, while loss of H3K9me3 enhances activation of transposable elements

In other species, H3K27me3 plays a crucial role in gene regulation, while H3K9me3 is involved in silencing of transposable elements [13,21,28]. Based on our observations from ChIP-seq data, we hypothesized that the two histone methylation marks have similar effects in *Z. tritici*. To test this hypothesis directly, we sequenced transcriptomes of two biological replicates of Zt09 and two independent transformants of the ∆*kmt1*, ∆*kmt6*, and Δ*k1/k6* deletion mutants after *in vitro* growth for 2 days representing exponential growth (Table S2).

First, we compared the total number of expressed genes. In total, 11,839 genes are annotated in the reference isolate [48]. Out of these, 8,906 are expressed (RPKM >2) in Zt09 during *in vitro* growth. The number of expressed genes is higher in both the Δ*kmt1* (9,259) and the Δ*k1/k6* (9,459) mutants, but to our surprise, lower in the Δ*kmt6* (8,717) mutants (Fig 2A, Table S3). This is in contrast to previous studies, where deletion of *kmt6* resulted in activation of otherwise silenced gene clusters and overall transcriptional activation [21,23,27,28]. We focused on differential gene expression between core and accessory chromosomes because genes on accessory chromosomes are silent under most conditions that have been tested. While 80 % of genes on core chromosomes are expressed in Zt09, only ~25% of genes located on accessory chromosomes display transcriptional activity. Transcription of genes on accessory chromosomes is higher in all mutant strains, ~40 – 50% (Fig 2A, Table S3), revealing gene activation on accessory chromosomes specifically upon removal of H3K27me3 or H3K9me3.

**Fig 2.**
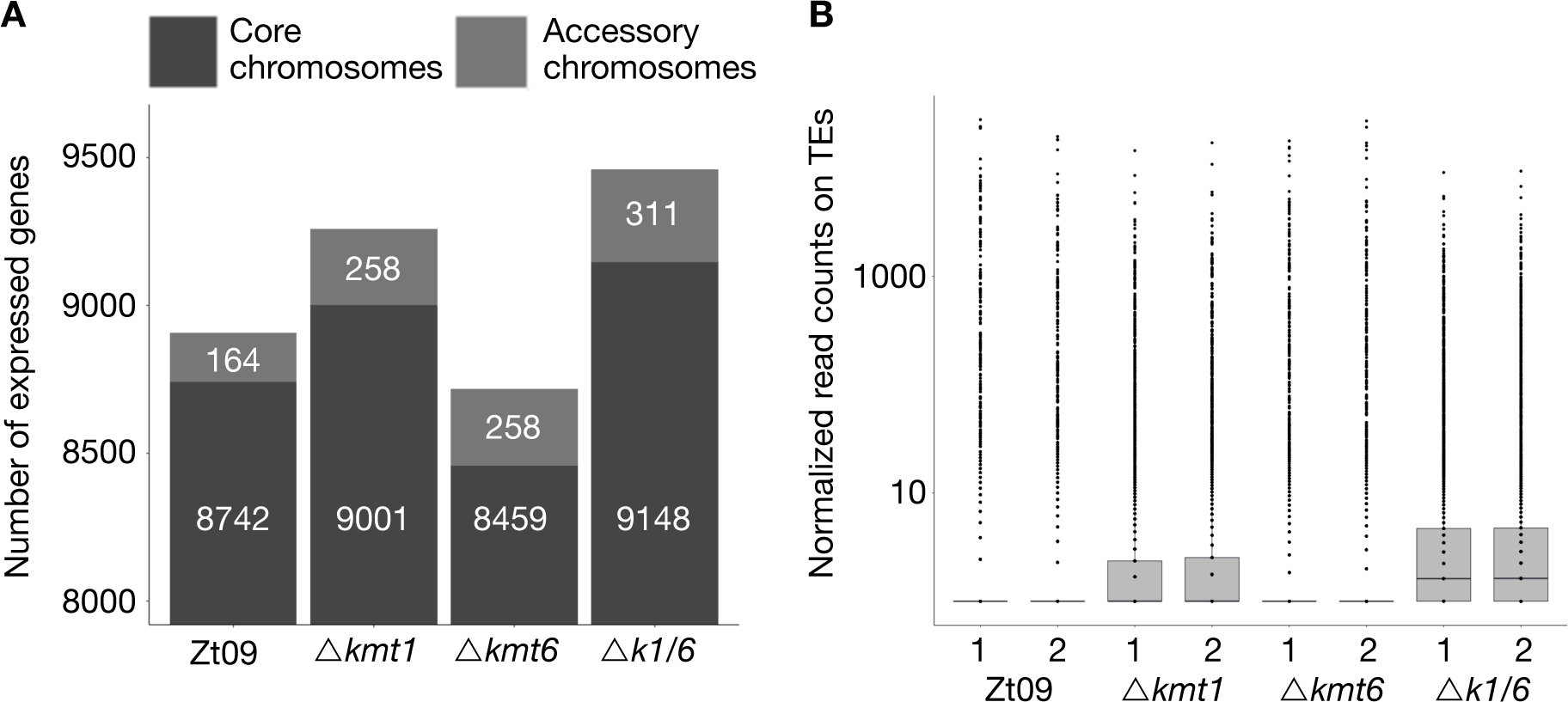
Gene (A) and transposon (B) expression increases in absence of H3K9me3, while loss of H3K27me3 alone decreases the number of expressed genes and does not impact transposon activity. **(A)** We compared the number of expressed genes in Zt09 and mutant strains. While in all mutants the number of expressed genes increases on accessory chromosomes, surprisingly loss of H3K27me3 alone in the Δ*kmt6* mutants resulted in a reduction of genes expressed on core chromosomes and only a small increase in numbers of genes expressed on accessory chromosomes. **(B)** Loss of H3K9me3, but not H3K27me3 alone, increases the number of transcripts originating from transposable elements. In absence of both marks (Δ*k1/k6*), the number further increases, likely because H3K27me3 moves to transposable elements in the Δ*kmt1* single mutant, facilitating silencing.

We further explored patterns of differential gene expression. Genome wide, 1,365 predicted genes were associated with H3K27me3 and 258 genes with H3K9me3 in Zt09. Interestingly, only a small fraction of genes associated with these histone marks were activated or differentially expressed in the mutants (Table S4). This indicates that loss of any of these methylation marks is not sufficient for transcriptional activation suggesting additional mechanisms involved in the transcriptional regulation of these genes.

In other fungi, removal of H3K9me3 and especially H3K27me3 was linked to the activation of certain gene classes, in particular secondary metabolite gene clusters [21,28,29]. To assess if genes with a specific function are enriched amongst the activated genes, we performed Gene Ontology (GO) enrichment analysis (topGO, Fisher’s exact test, *p*-value < 0.01). Consistent with the higher total number of expressed genes, we found the majority of differentially expressed (DE) genes (DESeq2, *P*_adj_ < 0.001, │log2 fold-change│ > 2) to be significantly upregulated in the Δ*kmt1* mutant (365 of 477) and in the Δ*k1/k6* mutant (368 of 477), whereas a majority of DE genes was downregulated in the Δ*kmt6* mutant (188 of 310) (Table S5).

We found two GO categories enriched amongst upregulated genes in Δ*kmt1* and Δ*k1/k6* mutants: DNA integration (GO:0015074) and RNA-dependent DNA replication (GO:0006278). Predicted functions assessed by BLAST analyses of the proteins encoded by the upregulated genes in these categories include reverse transcriptases, integrases, recombinases and genes containing transposon‐ or virus-related domains (Table S6). Consistent with these findings, we detected an increased number of transcripts originating from annotated transposable elements in the Δ*kmt1* and Δ*k1/k6* mutants, but not in Δ*kmt6* mutants (Fig 2B). This is in agreement with the strong association of transposable elements with H3K9me3 [31]. Transposons in subtelomeric regions and on accessory chromosomes show additional H3K27me3 enrichment. Removal of H3K9me3, but not of H3K27me3, appears to be responsible for transposon activation but transcription is further enhanced when both, H3K27me3 and H3K9me3 are removed in the *Δk1/k6* mutant (Fig 2B; Table S7).

Amongst the genes upregulated in the Δ*kmt6* mutant no GO categories were enriched but based on the previous finding of secondary metabolite activation, we further investigated possible roles of H3K9me3 and H3K27me3 in secondary metabolite gene regulation. Therefore, we identified putative secondary metabolite clusters in the *Z. tritici* reference genome using antiSMASH (antibiotics & Secondary Metabolite Analysis SHell) [51]. We found a total of 27 secondary metabolite clusters, all located on core chromosomes, and merged the identified genes with the existing gene annotation (Table S8). Except for the activation of one putative cluster on chromosome 7 in the Δ*k1/k6* mutant, we did not identify any differential expression of genes in secondary metabolite clusters. Based on these findings, we conclude that, unlike in other fungi, H3K9me3 and H3K27me3 are not involved in transcriptional regulation of secondary metabolites in *Z. tritici* under the tested conditions.

Taken together, removal of these histone modifications has little consequences for the expression of the vast majority of associated genes. As expected from its localization, loss of H3K9me3 increases expression of transposable elements while absence of H3K27me3 by itself has very little impact on transcriptional activation, thus suggesting that in this organism H3K27me3 does not delineate stereotypical “facultative heterochromatin”.

### Loss of H3K27me3 drastically reduces the loss of accessory chromosomes

Chromosome landmarks, namely centromeric and pericentric regions, telomere repeats and subtelomeric regions are similar on core and accessory chromosomes [31]. Accessory chromosomes are enriched with transposable elements but share the same TE families as core chromosomes [48]. Nevertheless, accessory chromosomes of *Z. tritici* are highly unstable, both during meiosis and vegetative growth *in vitro* and *in planta* [46,47]. The most striking feature that sets these chromosomes apart is almost chromosome-wide enrichment with H3K27me3 and, as a consequence of the higher TE content, increased enrichment with H3K9me3 [31]. To test whether loss of these modifications affects genome and chromosome stability in *Z. tritici*, we conducted two different long-term growth or “lab evolution” experiments to study genome stability and to detect dynamics of accessory chromosome losses in strains deficient for two important chromatin marks (Fig S8).

To assess whether the specific histone methylation pattern on accessory chromosomes contributes to instability of accessory chromosomes, we performed a short-term *in vitro* growth experiment over four weeks. Zt09, Δ*kmt6*, Δ*kmt1* and a Δ*k1/k6* double deletion mutant were used as progenitors. Each strain was grown in three replicate cultures and ~4% of the cell population was transferred to fresh medium every three to four days. After four weeks of growth, we plated dilutions of each culture to obtain single colonies that were subsequently screened by a PCR assay for the presence of accessory chromosomes (Table 2).

**Table 2.** Chromosome loss rates and frequency of individual accessory chromosome losses in the Zt09^1^ reference strain and mutants during short-term evolution experiments. Three replicate cultures were tested per strain.

| **Chromosome** | **Zt09** | ***Δkmt1*** | ***Δkmt6*** | ***Δk1/k6*** |
| --- | --- | --- | --- | --- |
| **14** | 18 | 26 | 2 | 13 |
| **15** | 8 | 0 | 0 | 0 |
| **16** | 9 | 2 | 0 | 2 |
| **17** | 0 | 0 | 5 | 0 |
| **19** | 0 | 0 | 0 | 3 |
| **20** | 2 | 51 | 2 | 17 |
| **21** | 1 | 6 | 1 | 6 |
| **total loss** | 38 | 85 | 10 | 41 |
| **total tested** | 576 | 576 | 576 | 576 |
| **loss rate (%)** | 6.6 | 14.8 | 1.7 | 7.1 |

<sup>1</sup> Strain Zt09 had previously spontaneously lost chromosome 18 [41].

Previously, we showed that accessory chromosomes are lost at a rate of ~7% in Zt09 and we documented that accessory chromosomes 14, 15 and 16 are more frequently lost than others [46]. Here we demonstrate that, in comparison to Zt09, the Δ*kmt1* mutant showed a significantly increased chromosome loss rate (one sided Fisher’s exact test for count data, *p*-value = 2.7 × 10^−6^). Interestingly, this was not due to an overall increase of accessory chromosome loss, but rather by the dramatically increased (*p*-value = 3.7 × 10^−9^) frequency of loss for chromosome 20 (Table 2). The chromosome loss rate of the other accessory chromosomes was either comparable to Zt09 (Chr. 14, 17, 19, 21) or even significantly lower (Chr. 15 and 16, *p*-values = 1.2 × 10^−3^ and 2.5 × 10^−5^). This suggests a special role of H3K9me3 for the maintenance of chromosome 20.

In contrast to the Δ*kmt1* mutants, we detected significantly fewer chromosome losses (*p*-value = 1.2 × 10^−4^) in the Δ*kmt6* mutants. Out of 576 tested colonies, only ten had lost an accessory chromosome. This represents a four times lower chromosome loss rate compared to wild type. Therefore, absence of H3K27me3 appears to promote stability of accessory chromosomes. Interestingly, chromosome 17 was lost with the highest frequency in this mutant (5/10) but was not lost in any of the other mutant strains or in the wild type.

The double deletion mutant displayed a similar chromosome loss rate as wild type but showed a chromosome loss distribution comparable to the Δ*kmt1* deletion strain with chromosome 20 being lost significantly more often (*p*-value = 1.23 10^−4^), and chromosomes 15 and 16 lost less frequently (*p*-values = 1.95 × 10^−3^ and 9 × 10^−3^, respectively). This suggests that the increase in chromosome stability in Δ*k1/k*6 compared to Δ*kmt1* is due to the removal of the destabilizing H3K27me3.

In summary, we found that loss of H3K27me3 increases accessory chromosome stability, suggesting a mechanistic explanation for how the widespread H3K27me3 enrichment on accessory chromosomes in normal cells contributes to the previously observed extraordinary chromosome instability.

### Loss of H3K9me3 promotes large-scale structural rearrangements mediated by TE instability and redistribution of H3K27me3

In a second evolution experiment, we addressed overall genome stability over a longer period of mitotic growth. The single mutants (Δ*kmt1* and Δ*kmt6*) and Zt09 were grown in triplicate cultures for ~6 months. We sequenced full genomes of progenitors and the evolved populations after 50 transfers to identify structural variations that arose during the experiment. All strains were illumina sequenced with ~100x coverage and paired-end reads were mapped to the reference genome of IPO323 and normalized to 1x coverage for visualization [40].

We focused our analysis on large scale chromosomal rearrangements such as duplications, deletions, and translocations. Structural variation was detected computationally from sequence alignments, validated experimentally by PFGE and Southern blotting, and additional rearrangements were identified by manual screening of mapped reads. Analysis of progenitor genomes revealed, except for the already known absence of chromosome 18 [41] and the previously described variations (point mutations and short indels) in Zt09 compared to the IPO323 reference genome [46], lower sequence coverage (~0.6x) on chromosome 17 in the Δ*kmt6* progenitor strain (Fig 3A). This difference can only be explained by a lower copy number in the sequenced pool of cells, suggesting loss of chromosome 17 in ~40% of the sequenced Δ*kmt6* cells, a chromosome loss that likely occurred at the very beginning of the experiment.

**Fig 3.**
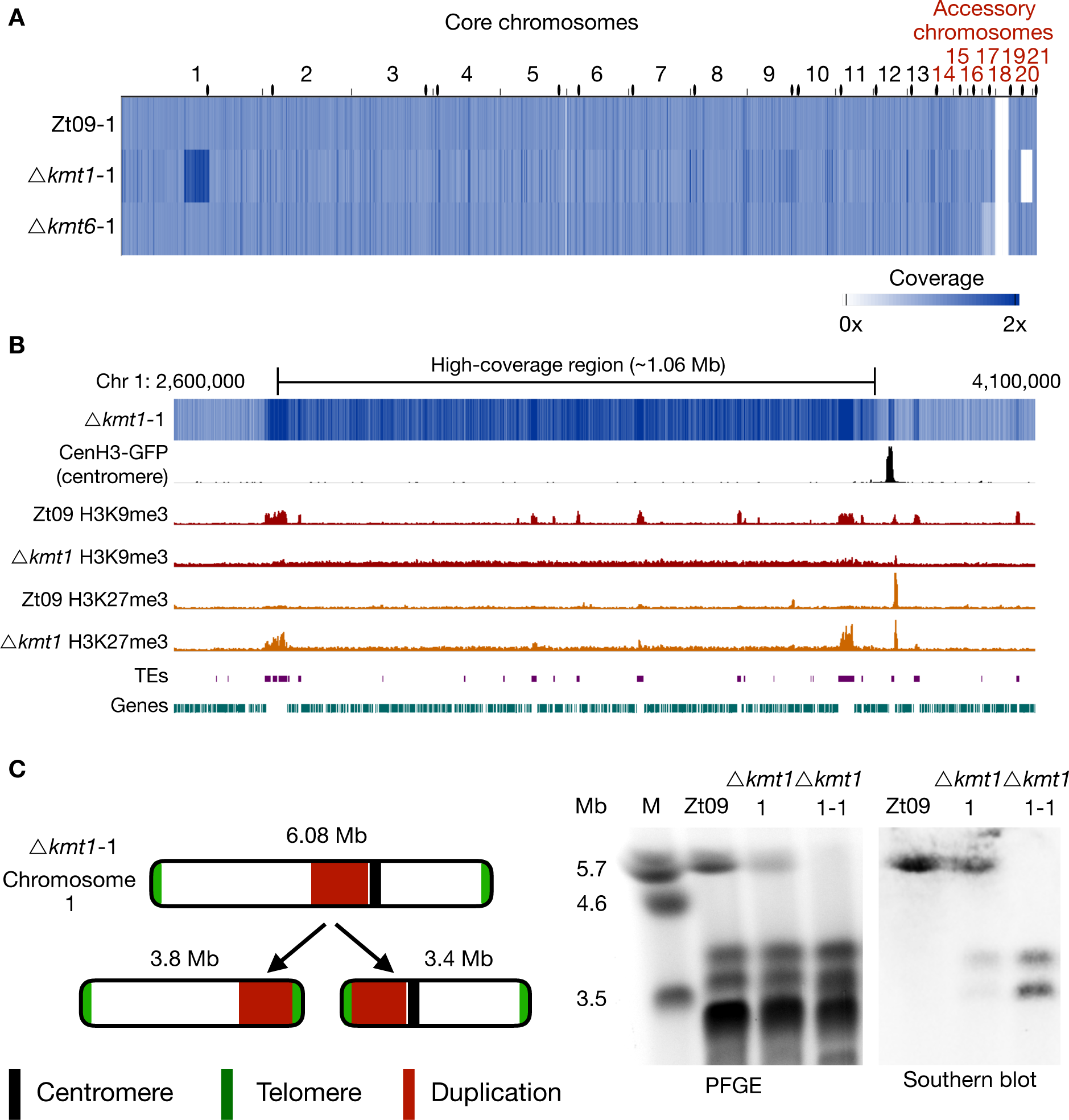
Genome sequencing of progenitor strains for the long-term growth experiment and analysis of structural variation in the *Δkmt1* progenitor. **(A)** The genomes of progenitor strains were sequenced and reads were mapped to the reference genome. Genome coverage was normalized to 1x coverage to allow identification and comparison of differences within and between strains. All strains are missing chromosome 18, as expected [41]. Δ*kmt6* has lower coverage (0.4x) of chromosome 17. *Δkmt1* lost chromosome 20 and, most notably, shows a long segment (~ 1Mb) of high-coverage (1.6x) on chromosome 1. Centromeres are indicated as black dots. **(B)** Examination of the high-coverage region breakpoints on chromosome 1. The first breakpoint locates within a TE-rich region that is enriched with H3K9me3 in Zt09 and shows new enrichment with H3K27me3 in *Δkmt1*. The second breakpoint is within a gene-rich region in close proximity to relocalized H3K27me3 and very close to the centromere (~15 kb). **(C)** Further analysis of this high-coverage region revealed *de novo* telomere formation at the breakpoints indicating a chromosome breakage at both ends of the high-coverage region. To validate chromosome breakage and possible new chromosome formation, we conducted PFGE and separated the large chromosomes of Zt09, of the *Δkmt1* progenitor strain (*Δkmt1*-1) and of a single clone originating from the *Δkmt1* progenitor strain stock (*Δkmt1*-1-1). Chromosome 1 (~6 Mb) is present in Zt09 and *Δkmt1*-1 (faint band), but not in the *Δkmt1*-1-1 single clone. We conducted Southern analysis on the PFGE blot using a sequence of the high-coverage region as a probe. It hybridized to the original chromosome 1 band in Zt09 and *Δkmt1*-1, but additionally to a ~3.4 Mb and ~3.8 Mb band in *Δkmt1*-1 and only to these bands in *Δkmt1*-1-1. This confirms the formation of new chromosomes, both containing the high-coverage region in some cells of the progenitor strain population.

Unexpectedly, the Δ*kmt1* progenitor displayed a long high-coverage (~1.6x) region on chromosome 1, suggesting that the region had been duplicated in ~60% of the sequenced Δ*kmt1* cells. Furthermore, this genome has a shorter chromosome 6 and does not contain chromosome 20 (Fig 3A and B, Table S9). The presence of this kind of structural variation in the progenitor strain is indicative for a high degree of genome instability in absence of Kmt1. Analysis of discordant reads mapped to both ends of the ~1 Mb high-coverage region on chromosome 1 revealed telomeric repeats (TTAGGG_n_), suggesting the formation of *de novo* telomeres. Pulsed-field gel electrophoresis (PFGE) and Southern analyses confirmed the formation of two new independent chromosomes both containing the high-coverage region and either the right or left arm of chromosome 1 (Fig 3C). The breakpoint on the left side coincides with a large TE-rich region that is associated with H3K9me3 in the wild type. Both breakpoints coincide with or are in close proximity to regions that show enrichment of relocated H3K27me3 in the *Δkmt1* mutant (Fig 3B), suggesting a possible link between relocated H3K27me3 and genome instability.

After six months of vegetative growth, we sequenced the pooled genomes of all nine ‘evolved’ populations. We found no evidence for large-scale genomic rearrangements in any of the evolved Zt09 or Δ*kmt6* populations (Fig 4A). Apart from seven small deletions or duplications (Table S9), the largest structural variation found in one of the evolved Δ*kmt6* populations (Δ*kmt6* 50-2), was a partial loss (~18 kb) at the right end of chromosome 15. However, we found variation in the read coverage of accessory chromosomes in all sequenced genomes indicating whole chromosome losses in individual cells of the population. The distinct dynamics of individual accessory chromosome losses were described in the previous section as part of the short-term growth results.

**Fig 4.**
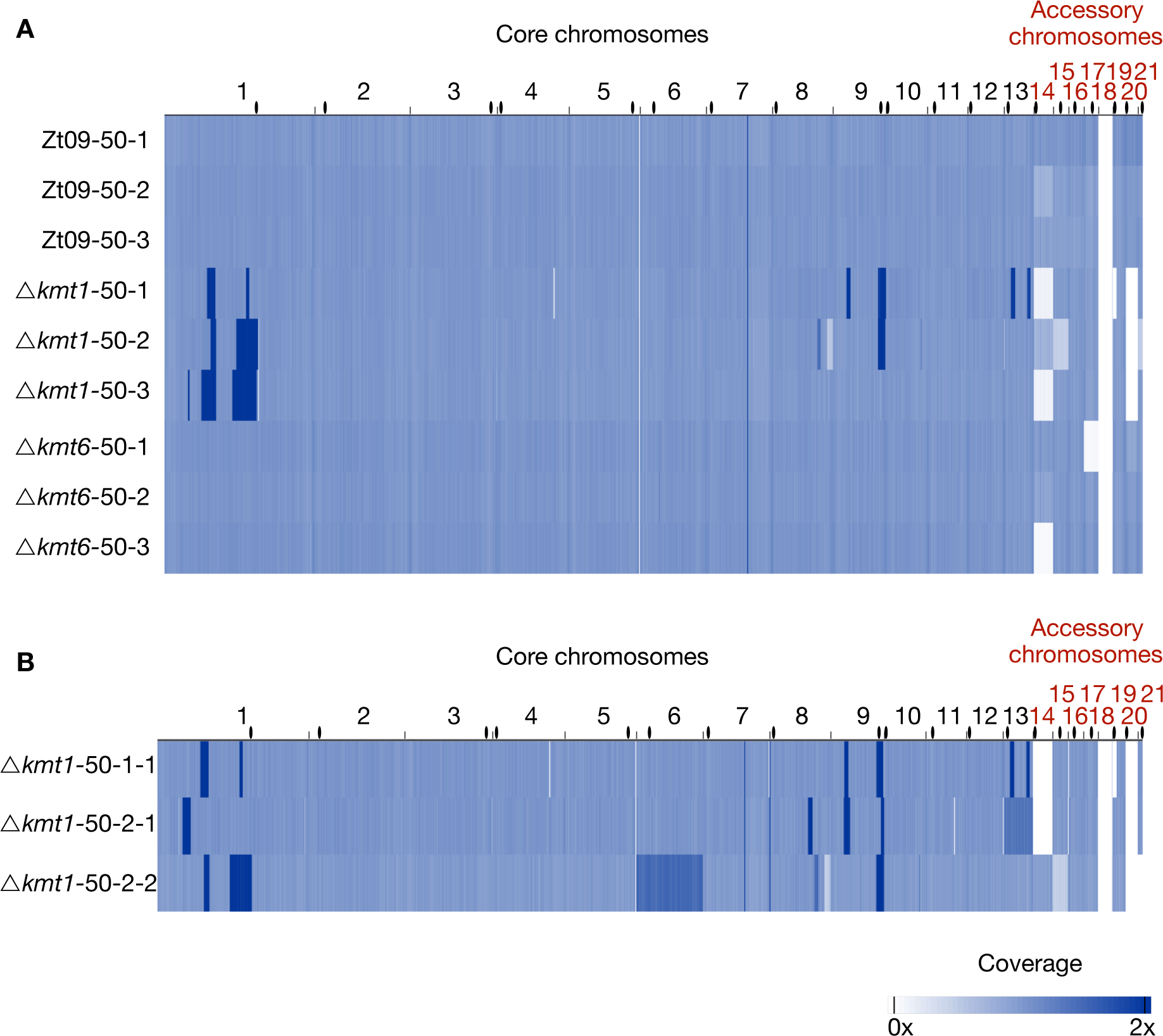
Genome sequencing of evolved populations and single clones originating from the long-term growth experiment. **(A)** Genomes of each replicate population after 50 transfers were sequenced and mapped to the reference. Coverage is normalized to 1x. Except for coverage differences on the accessory chromosomes, there are no large structural variations detectable for the evolved Zt09 and *Δkmt6* populations. In contrast, *Δkmt1* populations contain multiple high-coverage regions (dark blue) on core chromosomes as well as large deletions indicated by low (light blue) or no (white) coverage. **(B)** To further characterize structural variation in the evolved *Δkmt1* strains, three single clones originating from populations *Δkmt1*-50-1 (*Δkmt1*-50-1-1) and *Δkmt1*-50-2 (*Δkmt1*-50-2-1 and *Δkmt1*-50-2-2) were sequenced. Clones *Δkmt1*-50-1-1 and *Δkmt1*-50-2-2 show a very similar pattern as their respective populations, while *Δkmt1*-50-2-1 resembles a genotype that appears to be rare in population *Δkmt1*-50-2. High coverage on entire core chromosomes 13 (*Δkmt1*-50-2-1, 1.3x coverage) and 6 (*Δkmt1*-50-2-2, 1.5x coverage) indicates whole core chromosome duplications that are maintained in some nuclei. Centromeres are indicated as black dots.

In contrast to the few variations detected in the Zt09 and Δ*kmt6* populations, we found numerous large-scale high-coverage regions on different core chromosomes, chromosome breakages followed by *de novo* telomere formation, chromosomal fusions, as well as several smaller deletions and duplications in the evolved Δ*kmt1* populations (Fig 4A, Table S9). All three evolved Δ*kmt1* populations have large duplicated regions on chromosome 1 (Fig 5), but their locations as well as the resulting structural variations differ from the one identified in the progenitor strain (Fig 5A). This can be explained by independent events, as not all Δ*kmt1* progenitor cells underwent the rearrangement of chromosome 1 (Fig 3C), or by continuous structural rearrangement events as a consequence of the presence of large duplicated regions in the genome. Analyses of the affected regions and breakpoints indicate a connection between the structural variations of progenitor (compared to the reference) and evolved strains. In all evolved Δ*kmt1* populations, duplicated regions fully or partially overlap with the high-coverage region of the progenitor strain (Fig 5A-D).

**Fig 5.**
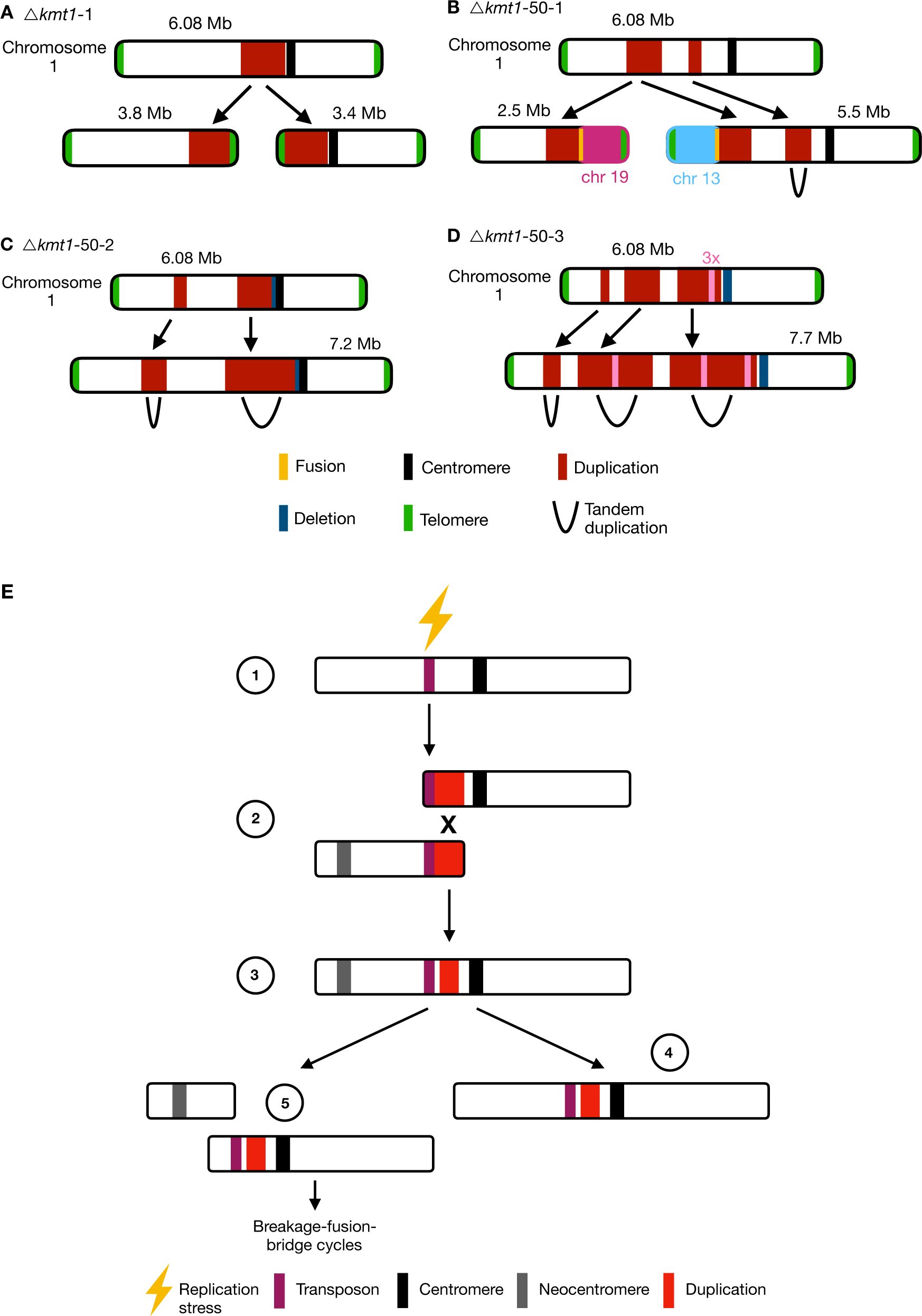
Different outcomes of structural variation of chromosome 1 in evolved *Δkmt1* strains. Upper panels display mapping to the reference genome including duplication indicated by higher read coverage (red), while the respective lower panels show structural rearrangements predicted by our structural variant analysis. **(A)** In the progenitor strain, a duplicated region was involved in the formation of two new chromosomes. At both termini of the duplication the chromosome broke and telomeric repeats were added *de novo* to these breakpoints (see Fig. 3). Thus, two new chromosomes were formed, both containing the duplicated sequence. This structural variation is not found in all cells in the Δ*kmt1* progenitor strain and the structural variation that arose in the evolved strains **(B-D)** can therefore be the result of rearrangements of the reference chromosome 1 or the two newly formed chromosomes. **(B)** In the evolved population Δ*kmt1*-50-1, two duplicated sequences were detected. The borders of the first region mark chromosome breakages that are fused to telomeres of other chromosomes. The first breakpoint is attached to the telomere of chromosome 13 forming a new 5.5 Mb chromosome while the second breakpoint is fused to the telomere of chromosome 19 (new 2.5 Mb chromosome). The second duplicated region represents a tandem duplication located on the new 5.5 Mb chromosome that falls within the duplicated region of the progenitor strain. **(C)** Population Δ*kmt1*-50-2 contains two duplicated regions, that both resemble tandem duplications. The second duplication is very similar to the one found in the progenitor strain but includes half of the centromere and has a deletion, where the breakpoint close to the centromere in the progenitor strain is located. **(D)** Population Δ*kmt1*-50-3 displays three duplicated sequences that all form tandem duplications resulting in the formation of a 7.7 Mb version of chromosome 1. The third duplicated region is, as in population Δ*kmt1*-50-2, very similar to the one in the progenitor strain. However, in this case the complete centromere-associated sequence is deleted. Furthermore, a ~50 kb region inside the third duplicated region exhibits 3x sequencing coverage and is found in between the tandem duplication of the second duplicated region (see Table S9). **(E) Model for the formation of large structural rearrangements in the *Δkmt1* mutants over time. 1:** Replication stress at repeated sequences enriched with relocalized H3K27me3 promotes structural variation in form of deletion or duplication. **2:** Large segmental duplications arise during that process that are followed by chromosome breakage and new chromosomes are formed by adding *de novo* telomeric repeats at chromosomal breakpoints. While one of the chromosomal parts contains the original centromere, the other *de novo* chromosome forms a neocentromere. **3:** The duplicated sequences are targets for mitotic recombination resulting in chromosome fusion. The chromosome is now dicentric. **4:** To stabilize the chromosome, one of the two centromeres is inactivated, either epigenetically or by deletion of the underlying sequence. **5:** Alternatively, the dicentric chromosome becomes unstable during mitosis and breaks between the two centromeres. The broken chromosome ends are repaired either by *de novo* telomere formation or fusion to a different chromosome, giving rise to new breakage-fusion-bridge cycles in following rounds of mitotic cell divisions.

Since populations reflect a mixture of distinct genotypes, we also sequenced three single Δ*kmt1* clones originating from the populations from transfer 50 to characterize the structural variation in more detail (Fig 4B). The single clones were selected based on different PFGE karyotypes (Fig S9) and originated from population *Δkmt1-*50-1 (Δ*kmt1-*50-1-1) and Δ*kmt1-*50-2 (Δ*kmt1-*50-2-1 and Δ*kmt1-*50-2-2). As two of these single clones (Δ*kmt1-*50-1-1 and Δ*kmt1-*50-2-2) largely resemble the genotypes found in their respective populations, we conclude the presence of a predominant genotype in each evolved replicate population. However, Δ*kmt1-*50-2-1 clearly differs from this genotype and therefore reveals the existence of additional, rarer genotypes in populations. Relatively small deletions and duplications (up to 30 kb) as well as chromosome breakage followed by *de novo* telomere formation were found on almost all chromosomes. These occurred mainly linked to annotated transposable elements (Table S10) whereby loss of H3K9me3 likely promoted instability. However, major rearrangements, including chromosomal fusions, were always linked to large segmental duplications (Fig S10). In two strains we detected higher coverage of entire core chromosomes indicating core chromosome duplications (Fig 4B). Results from read coverage and PCR analyses indicate that Δ*kmt1-*50-2-2, as well as the majority of the Δ*kmt1-*50-2 population, may have undergone a whole genome duplication.

To investigate whether the underlying sequence is involved in the formation of large-scale rearrangements, we analyzed the breakpoints of each duplicated region. The location of breakpoints does not show a clear TE-associated pattern as observed for the smaller deletions or chromosome breakages. Out of 28 analyzed breakpoints, only seven are directly located within annotated transposable elements, while thirteen fall into genes, seven are intergenic and one is located in the centromere (Table S11). Considering all structural rearrangements in the three sequenced single clones, we found that out of 62 events, 34 were associated (direct overlap or <5 kb distance) to regions that show enrichment for H3K27me3 (Table S12). Based on these observations, we hypothesize that two non-exclusive pathways, namely TE-associated instability caused by loss of H3K9me3 or invasion of H3K27me3 with increased recombination activity, mayserve as initial events, which are followed by continuous rearrangements resulting in a spectrum of structural variation (Fig 5E).

## Discussion

### H3K27me3 destabilizes accessory chromosomes

We investigated the effects of loss of two important heterochromatin associated histone modifications, H3K9me3 and H3K27me3, on chromatin organization, transcription and genome stability and characterized phenotypes of the deletion mutants. Loss of H3K9me3 allows relocalization of H3K27me3 in *kmt1* deletion mutants, which has great impact on genome and chromosome stability, resulting in numerous large-scale rearrangements. In contrast, the genomes of evolved Δ*kmt6* and Zt09 strains revealed only few and relatively minor changes. Unexpectedly, the presence of H3K27me3 impacts chromosome stability by either destabilizing whole chromosomes in normal cells, supported by the high loss-rate in the reference strain compared to the Δ*kmt6* mutants, or by mislocalization as shown by the increased sequence instability in the Δ*kmt1* mutants. Taken together, enrichment with H3K27me3 in wild type cells is a main driver of mitotic chromosome instability.

We propose different scenarios for how chromosomes may get lost during mitosis and how H3K27me3 may be linked to these processes. For example, accessory chromosomes may not be accurately replicated whereby only one sister chromatid is transmitted. Alternatively, non-disjunction of sister chromatids during mitosis produces one cell with two copies and one cell lacking the respective chromosome. Previous microscopic studies of *Z. tritici* expressing GFP-tagged CENPA/CenH3 proteins indicated that core and accessory chromosomes might be physically separated in the nucleus [31]. Interestingly, previous studies showed that H3K27me3-enriched chromatin localizes close to the nuclear periphery and loss of H3K27me3 enables movement of the previously associated regions to the inner nucleus in mammals and fungi [52,53]. Proximity to the nuclear membrane and heterochromatic structure can furthermore result in differential, and often late, replication timing [54,55]. Loss of H3K27me3 and the associated movement to the inner nuclear matrix might therefore alter replication dynamics of accessory chromosomes resulting in higher rates of faithfully replicated chromosomes and lower rates of mitotic losses.

Heterochromatic regions, especially associated with H3K27me3, tend to cluster together and form distinct foci in the nucleus of *Drosophila melanogaster* visualized by microscopic analyses [56,57], and loss of H3K27me3 reduces interaction between these regions [58]. We hypothesize that enrichment of H3K27me3 on the entire accessory chromosomes maintains physical interactions that persist throughout mitosis. This may decrease the efficiency of separation of sister chromatids resulting in loss of the chromosome in one cell and a duplication in the other cell. So far, we have focused our screening on chromosome losses but determining the exact rates of accessory chromosome duplications is necessary to test this hypothesis. Genome sequencing of *Z. tritici* chromosome loss strains revealed that duplications of accessory chromosomes can occur [46]. Similarly, B chromosomes in rye are preferentially inherited during meiosis by non-disjunction of sister chromatids during the first pollen mitosis [59], indicating that deviation from normal chromosome segregation occurs. Accessory chromosomes are commonly found in natural isolates of *Z. tritici*, despite the high loss rates we demonstrated during mitotic growth [46]. This observation implies the presence of other mechanisms that counteract the frequent losses of accessory chromosomes. Recent analyses of meiotic transmission showed that unpaired accessory chromosomes are transmitted at higher rates in a uniparental way [60,61]. We propose that H3K27me3 is involved in accessory chromosome instability and transmission both during mitosis and meiosis.

### H3K9me3 loss allows invasion by H3K27me3 and results in genome instability

Loss of H3K9me3 had severe effects on genome stability and growth and reproduction *in vitro* and *in planta*, while loss of H3K27me3 only resulted in minor differences to wild type growth and, unexpectedly, rather promoted than decreased genome stability. In contrast to the experimentally evolved Zt09 and Δ*kmt6* mutants, we detected a high number of smaller (up to 30 kb) deletions and duplications, chromosome breakages and several gross chromosomal rearrangements linked to large duplications in the Δ*kmt1* mutants. Absence of H3K9me2/3 has been associated with chromosome and genome instability in other organisms [13,14,62,63]. Smaller deletions, duplications and chromosome breakages resulting in shortened chromosomes due to loss of chromosome ends that we identified in the Δ*kmt1* mutants, correlate with transposable elements, enriched with H3K9me3 in wild type. Replication of heterochromatin-associated DNA is challenging for the cell as repeated sequences can form secondary structures that can stall the replication machinery [64]. Instability of repeated sequences has consequently been linked to errors during DNA replication [65,66]. Normally, the replication machinery and heterochromatin-associated proteins work together to ensure faithful replication and genome integrity [67]. In *Caenorhabditis elegans*, loss of H3K9me promotes transposable element transcription and formation of R-loops (RNA:DNA hybrids) at repeated sequences during replication resulting in copy-number variations [13]. This phenomenon may explain the accumulation of small deletions and duplications, chromosome breakage or the formation of large segmental duplications that we observed in the Δ*kmt1* mutants (Tables S8 and S9). Furthermore, the structural variation that arises depends on the mode of DNA repair following the DNA damage. Double-strand breaks can be repaired by non-homologous end joining causing deletions or translocations or by homologous recombination [68]. Alternatively, they can be healed by generation of telomeric repeats and *de novo* telomere formation [69]. The structural rearrangements detected in the Δ*kmt1* mutants indicate that repair of double-strand breaks involves both non-homologous end joining and *de novo* telomere formation. We propose that the main factor for genome instability is replication-associated instability of repeated sequences subsequently promoting the formation of large-scale rearrangements (Fig 5E).

Not all breakpoints of rearrangements, especially of the large duplicated sequences, were associated with transposable elements, however. We found that duplicated sequences in the experimentally evolved Δ*kmt1* mutants fully or partially overlap with the duplicated regions of the Δ*kmt1* progenitor strain. This strongly indicates that structural variations are subject of continuous rearrangements, resulting in rearrangements that are not directly linked to the initial event. It is important to note that the rearrangements and genotypes we detected are the result of selection during our long-term growth experiments and thus do not necessarily reflect the full spectrum of rearrangements occurring in the *Δkmt1* mutants; many additional structural variants may have disappeared quickly from the population or even included lethal events.

Concomitant with loss of H3K9me3 in the Δ*kmt1* strains, we found relocalization of H3K27me3 to former H3K9me3 regions. A similar redistribution of H3K27me3 in absence of heterochromatin factors has been reported in plants and animals [70–72] and other fungi [22,27,73]. In *N. crassa*, redistribution of H3K27me3 in a Δ*kmt1* (*dim-5*) mutant background results in severe growth defects and increased sensitivity to genotoxic stress that can be rescued by elimination of H3K27me3, indicating that aberrant H3K27me3 distribution severely impacts cell viability [27]. Although we did not see that phenotypic defects *in planta* or in the *in vitro* stress assay are rescued in the Δ*k1/k6* double mutants, the chromosome-loss rate was reduced compared to *Δkmt1* mutants indicating a stabilizing effect when H3K27me3 is absent. We found that some breakpoints of the rearrangements in the Δ*kmt1* mutants do not only coincide with regions that have lost H3K9me3 enrichment but that also show enrichment with the invading H3K27me3. This raises the question whether sequences associated with H3K27me3 are more susceptible to genome instability. Regions enriched with H3K27me3 have been shown to exhibit a high degree of genetic variability in form of mutations, increased recombination or structural variation compared to the rest of the genome [21,23,30,31,74,75]. Experimental evolution in *Fusarium fujikuroi* showed that increased H3K27me3 levels in subtelomeric regions coincided with increased instability [76] and we previously detected a highly increased rate of chromosomal breakage under stress conditions in subtelomeric, H3K27me3 regions in *Z. tritici* [46]. These observations together with our findings strongly indicate that H3K27me3 plays a pivotal role in decreasing genome stability.

### Evolutionary implications of chromatin and genome instability

We found several large-scale genome rearrangements including new chromosomes containing very large duplicated regions and chromosomal translocations in the evolved *Δkmt1* mutants. The formation of new chromosomes described in this study suggests a mechanistic basis for the emergence of accessory chromosomes in *Z. tritici*. The newly generated chromosomes contain a relatively high proportion of duplicated sequences, but functional centromeres and telomeres are readily maintained. Accessory chromosomes in *Z. tritici* do not, however, contain a high number of paralogs of core chromosome genes [41]. Duplicated genes may not be functional over long evolutionary timescales, because they can become pseudogenes via mutational drift, thus resulting in the more typical gene sparse, heterochromatic accessory chromosomes [30]. Indeed, the origin of accessory chromosomes from core chromosomes has been proposed for fungal chromosomes [77,78] and shown in several plant species [79]. In future studies, we will therefore investigate the mechanisms for *de novo* centromere and telomere formation that must occur to generate additional chromosomes, and we will follow the fate of segmental duplications that were generated in this study.

Furthermore, chromosomal rearrangements initiated by translocations or mitotic recombination can result in the formation of dicentric chromosomes that are instable during mitosis giving rise to new chromosome rearrangements mediated by breakage-fusion-bridge cycles (Fig 5E) [80,81]. A previous study demonstrated the formation of a new accessory chromosome by breakage-fusion-bridge cycles during meiosis in *Z. tritici* [82]. In our study, we find evidence for the occurrence of breakage-fusion-bridge cycles by the observed fusion events between chromosomes 1 and 19 and between 1 and 13 (Fig S10). For these chromosomes we observe that the breakpoints of a large duplicated sequence on chromosome 1 are fused to telomeres of chromosomes 13 and 19. Both chromosomes broke close to their centromeres resulting in loss of the centromeric sequence. A mechanism to avoid breakage-fusion-bridge cycles is the inactivation of centromeres. This can either be accomplished by epigenetic inactivation or by deletion of the underlying sequence [83]. We detected deletions and partial duplications of centromeric DNA (Table S9, Fig 5E) but further analyses to map localization of CENPA/CenH3 must be conducted to investigate neocentromere formation and centromere inactivation. As we found evidence that new chromosomes without the original centromere can be generated (Fig 3), we hypothesize that neocentromeres are readily established on these chromosomes. Previously, we have shown that the structure of *Z. tritici* centromeres is unusual, as they do not display typical pericentric heterochromatin regions but contain actively transcribed genes [31]. These characteristics are similar to neocentromeres found in *Candida albicans* [84,85] suggesting that centromere formation in *Z. tritici* is highly dynamic.

The presence of Kmt1 and of H3K9me3 respectively, is essential to maintain genome integrity in this fungus. TE-mediated rearrangements may be involved in the genetic variability detected in *Z. tritici* isolates [86–88] and have been suggested as drivers of genome evolution in various species [89–91]. Our findings concerning the role of H3K9me3 for genome stability provide a basis for future studies focusing on the influence of heterochromatin on structural genome rearrangements using *Z. tritici* as a model organism.

We found that, unlike for H3K9me3, presence and not absence of H3K27me3 is linked to genome instability. Surprisingly, loss of H3K27me3 does not result in dramatic changes of overt phenotypes and is also not clearly linked to transcriptional activation in *Z. tritici*. This allowed us to uncouple the transcriptional and regulatory effects of H3K27me3 from the influence on chromatin stability and will in the future result in further mechanistic insights on the influence of histone modifications on chromosome stability.

## Materials and Methods

### Culturing conditions of fungal and bacterial strains

*Zymoseptoria tritici* strains were cultivated on solid (2% [w/v] bacto agar) or in liquid YMS (yeast-malt-sucrose) medium (0.4% [w/v] yeast extract, 0.4% [w/v] malt extract, 0.4% [w/v] sucrose per 1 L). Liquid cultures were inoculated from plate or directly from glycerol stocks and grown for 3 – 4 days at 18°C in a shaking incubator at 200 rpm. Plates were inoculated from glycerol stocks and grown for 5 – 6 days at 18°C. *Escherichia coli* TOP10 cells were grown overnight in dYT (1.6% [w/v] tryptone, 1% [w/v] yeast extract, 0.5% [w/v] NaCl and 2% bacto agar for solid medium) supplemented with antibiotics for plasmid selection (40 µg/mL kanamycin) at 37°C and at 200 rpm for liquid cultures. *A. tumefaciens* strain AGL1 was grown in dYT containing rifampicin (50 µg/mL) and carbenicillin (100 µg/mL) supplemented with antibiotics for plasmid selection (40 µg/mL kanamycin) at 28°C at 200 rpm in liquid culture for 18 h and on plate at 28°C for two days.

### Transformation of *Z. tritici*

*Z. tritici* deletion and complementation strains were engineered using *A. tumefaciens*-mediated transformations as described before [49,92]. Flanking regions of the respective genes were used to facilitate homologous recombination for integration at the correct genomic location. The plasmid pES61 (a derivate of the binary vector pNOV-ABCD [49]) was used for targeted gene deletion and complementation. Plasmids were assembled using a restriction enzyme-based approach or Gibson assembly [93]. Plasmids were amplified in *E. coli* TOP10 cells and transformed in the *A. tumefaciens* strain AGL1 as described previously [94]. Gene deletions of *kmt1* (*Zt09_chr_1_01919*) and *kmt6* (*Zt09_chr_4_00551*) were facilitated by replacement of the respective ORF with a hygromycin resistance cassette. The *kmt1/kmt6* double deletion mutant was constructed by integrating a nourseothricin resistance cassette replacing *kmt1* in a *kmt6* deletion mutant background. Complementation constructs containing the respective gene and a G418 resistance cassette were integrated at the native loci in the deletion strains. All plasmids and strains constructed in this study are listed in Table S1. Transformed strains were screened by PCR for correct integrations of the construct followed by Southern blot [95] using DIG-labeled probes generated with the DIG labeling kit (Roche, Mannheim, Germany) following manufacturer’s instructions.

### DNA isolation for PCR screenings and Southern blotting

For rapid PCR screenings (candidates for transformation and chromosome loss), a single *Z. tritici* colony was resuspended in 50 µL 25 mM NaOH, incubated at 98°C for 10 min and afterwards 50 µL 40 mM Tris-HCl pH 5.5 were added. 4 µl mix subsequently was used as template for PCRs. For DNA extraction for Southern blotting, we used a standard phenol-chloroform extraction protocol [96] for DNA isolation.

### Phenotypic characterization *in vitro*

For the *in vitro* growth assays, liquid YMS cultures were inoculated with 100 cells/µL (OD_600_ = 0.01); cells were grown in 25 mL YMS at 18°C and 200 rpm. For each mutant and complementation strain, two transformants (biological replicates), and three replicate cultures per transformant (technical replicates) were used. For the reference strain Zt09, two separate pre-cultures were grown as biological replicates and each pre-culture was used to inoculate three replicate cultures. OD_600_ was measured at different time points throughout the experiment until the stationary phase was reached. The R package growthcurver [97] was used to fit the growth curve data enabling to compare *in vitro* growth of the different strains.

To test the tolerance of mutant and reference strains towards different stressors, we performed an *in vitro* stress assay on YMS plates. Each plate contained additives constituting different stress conditions. Cell suspensions containing 10^7^ cells/mL and a tenfold dilution series down to 100 cells/mL were prepared; 3 µL of each dilution were pipetted on solid YMS containing the following additives: 0.5 M NaCl, 1 M NaCl, 1 M sorbitol, 1.5 M sorbitol, 1.5 mM H_2_O_2_, 2 mM H_2_O_2_, 300 µg/mL Congo red, 0.01 % MMS (methyl methane sulfonate), 0.025 % MMS, 1 µg/mL actinomycin D and 1.5 µg/mL actinomycin D. Furthermore, we included a H_2_O-agar (2% bacto agar) plate. All plates were incubated at 18°C for six days, except for one YMS plate that was incubated at 28°C to test for thermal stress responses.

### Phenotypic characterization *in planta*

Seedlings of the susceptible wheat cultivar Obelisk (Wiersum Plantbreeding BV, Winschoten, The Netherlands) were potted (three plants per pot) after four days of pre-germination and grown for seven more days. Single cell suspensions of mutant and reference strain were prepared (10^8^ cells / mL in H_2_O with 0.1% Tween 20) and brush inoculated on a marked area of the second leaf. Following inoculation, the plants were incubated in sealed plastic bags containing ~ 1 L of H_2_O for 48 h providing high humidity to promote infections. Growth conditions for the plants throughout the complete growth phase and infection were 16 h light (200 µmol/m^−2^s^−1^) and 8 h dark at 20°C and 90% humidity. First appearances of symptoms, necrosis or pycnidia, were assessed by manual inspection of every treated leaf. 21 days post infection, inoculated leaves were finally screened for infection symptoms. Visual inspection of each leaf was performed to evaluate the percentage of leaf area covered by necrosis and pycnidia. Six different categories were differentiated based on the observed coverage (0: 0%, 1: 1 – 20%, 2: 21 – 40%, 3: 41 – 60%, 4: 61 −80%, 5: 81 – 100%). Furthermore, automated symptom evaluation was performed by analysis of scanned images of infected leaf areas as described previously [98].

### Long-term evolution experiment

For the long-term evolution experiment (~6 months), cells were inoculated directly from the glycerol stocks into 20 mL liquid YMS cultures. We used Zt09, Δ*kmt6* (#285) and Δ*kmt1* (#68), each strain grown in triplicates. Every three to four days, cells were transferred to new YMS medium. Cells were grown at 18°C and 200 rpm. For every transfer, cell density of the cultures was measured by OD_600_ and the new cultures were inoculated with a cell density of ~ 100 cells /µL (correlating to a transfer of 0.1% of the population). After 50 transfers, the genomes of the evolved populations and each progenitor strain were sequenced. Additionally, three genomes of single clones derived from the *Δkmt1* populations after 50 transfers were sequenced to characterize genome rearrangements in more detail.

### Short-term evolution experiment

For the short-term evolution experiment over a time period of four weeks, cultures were inoculated from single colonies grown on solid YMS. Zt09, *Δkmt6* (#285), *Δkmt1* (#80), and the *Δk1*/*Δ*k6 (#23) double mutant were grown in triplicate YMS cultures. For this experiment we used a different independent *Δkmt1* mutant clone (#80), as we discovered that the strain used in the previous long-term evolution experiment (#68) was missing chromosome 20. Every three to four days, 900 µL culture were transferred to 25 mL fresh YMS (correlating to a transfer of ~ 4% of the population). After four weeks of growth (including eight transfers to new medium) at 18°C and 200 rpm, cultures were diluted and plated on YMS agar to obtain single colonies. These single colonies were PCR screened for presence of accessory chromosomes as described in [46].

### Pulsed-field gel electrophoresis (PFGE)

Cells were grown in YMS medium for five days and harvested by centrifugation for 10 min at 3,500 rpm. We used 5 × 10^8^ cells for plug preparation that were washed twice with Tris-HCl, pH 7.5, resuspended in 1 mL TE buffer (pH 8) and mixed with 1 mL of 2.2% low range ultra agarose (Bio-Rad, Munich, Germany). The mixture was pipetted into plug casting molds and cooled for 1 h at 4°C. Plugs were placed to 50 mL screw cap Falcon tubes containing 5 mL of lysis buffer (1% SDS; 0.45 M EDTA; 1.5 mg/mL proteinase K[Roth, Karlsruhe, Germany]) and incubated for 48 h at 55°C while the buffer was replaced once after 24 h. Chromosomal plugs were washed three times for 20 min with 1 X TE buffer before storage in 0.5 M EDTA at 4°C. PFGE was performed with a CHEF-DR III pulsed-field electrophoresis system (BioRad, Munich, Germany). Separation of mid-size chromosomes was conducted with the settings: switching time 250 s – 1000 s, 3 V/cm, 106° angle, 1% pulsed-field agarose in 0.5 X TBE for 72 h. Large chromosomes were separated with the following settings: switching time 1000 s – 2000 s, 2 V/cm, 106° angle, 0.8% pulsed-field agarose in 1 X TAE for 96 h. *Saccharomyces cerevisiae* chromosomal DNA (BioRad, Munich, Germany) was used as size marker for the for mid-size chromosomes, *Schizosaccharomyces pombe* chromosomal DNA (BioRad, Munich, Germany) for the large chromosomes. Gels were stained in ethidium bromide staining solution (1 μg/mL ethidium bromide in H_2_O) for 30 min. Detection of chromosomal bands was performed with the GelDocTM XR+ system (Bio-Rad, Munich, Germany). Southern blotting was performed as described previously (Southern 1975) but using DIG-labeled probes generated with the PCR DIG labeling Mix (Roche, Mannheim, Germany) following the manufacturer’s instructions.

### ChIP-sequencing

Cells were grown in liquid YMS medium at 18°C for 2 days until an OD_600_ of ~ 1 was reached. Chromatin immunoprecipitation was performed as previously described [99] with minor modifications. We used antibodies against H3K4me2 (#07-030, Merck Millipore), H3K9me3 (#39161, Active Motif) and H3K27me3 (#39155, Active Motif). ChIP DNA was purified using SureBeads™ Protein G Magnetic Beads (Bio-Rad, Munich, Germany) and, replacing phenol/chloroform extractions, we used the ChIP DNA Clean & Concentrator Kit (Zymo Research, Freiburg, Germany). We sequenced two biological and one additional technical replicate for Zt09, Δ*kmt1*, Δ*kmt6*, and the Δ*k1*/*k6* strains. Sequencing was performed at the OSU Center for Genome Research and Biocomputing on an Illumina HiSeq2000 or HiSeq3000 to obtain 50-nt reads and at the Max Planck Genome Center, Cologne, Germany (https://mpgc.mpipz.mpg.de/home/) on an Illumina Hiseq3000 platform obtaining 150-nt reads (Table S2).

### RNA-sequencing

For RNA extraction, cells were grown in liquid YMS at 18°C and 200 rpm for two days until an OD_600_ of ~ 1 was reached. Cells were harvested by centrifugation and ground in liquid nitrogen. Total RNA was extracted using TRIzol (Invitrogen, Karlsruhe, Germany) according to manufacturer’s instructions. The extracted RNA was further DNAse-treated and cleaned up using the RNA Clean & Concentrator-25 Kit (Zymo Research, Freiburg, Germany). RNA samples of two biological replicates of Zt09, Δ*kmt1*, Δ*kmt6*, and the Δ*k1*/*k6* double mutant were sequenced. Poly(A)-captured, stranded library preparation and sequencing were performed by the Max Planck-Genome-centre Cologne, Germany (https://mpgc.mpipz.mpg.de/home/) on an Illumina Hiseq3000 platform obtaining ~ 20 million 150-nt reads per sample (Table S2).

### Genome sequencing

Genomic DNA for sequencing was prepared as described previously [100]. Library preparation and genome sequencing of the progenitor strains used for the evolution experiments were performed at Aros, Skejby, Denmark using an Illumina HiSeq2500 platform obtaining 100-nt paired-end reads. Library preparation (PCR-free) and sequencing of the evolved populations and the three evolved single *Δkmt1* mutants were performed by the Max Planck Genome Center, Cologne, Germany (https://mpgc.mpipz.mpg.de/home/) on an Illumina HiSeq3000 platform resulting in 150-nt paired-end reads (Table S2).

### Short read mapping and data analysis

A detailed list of all programs and commands used for mapping and sequencing data analyses can be found in the supplementary text S1. All sequencing data was quality filtered using the FastX toolkit ((http://hannonlab.cshl.edu/fastx_toolkit/) and Trimmomatic [101]. RNA-seq reads were mapped using hisat2 [102], mapping of ChIP and genome data was performed with Bowtie2 [103]. Conversion of sam to bam format, sorting and indexing of read alignments was done with samtools [104].

To detect enriched regions in the ChIP mappings, we used HOMER [105]. Peaks were called individually for replicates and merged with bedtools [106]. Only enriched regions found in all replicates were considered for further analyses. Genome coverage of enriched regions and overlap to genes and transposable elements was calculated using bedtools [106].

We used cuffdiff [107] to calculate RPKM values and to estimate expression in the different strains. Raw reads mapping on genes and transposable elements were counted by HTSeq [108], differential expression analysis was performed in R [109] with DESeq2 [110]. Cutoff for significantly differentially expressed genes was padj < 0.001 and |log2 fold-change| > 2. The R package topGO [111] was used to perform gene ontology enrichment analyses. Fisher’s exact test (*p*-value < 0.01) was applied to detect significantly enriched terms in the category ‘biological process’.

To detect structural variation in the sequenced genomes, we used SpeedSeq [112] and LUMPY [113]. All detected variation was further verified by manual visual inspection. Visualization was performed with the integrative genome browser (IGV) [114].

### Data availability

Sequencing raw reads (FASTQ files) of all genomic, ChIP-seq and RNA-seq data are available online at Sequence Read Archive (SRA) under BioProject ID PRJNA494102. Strains are available upon request.

## Acknowledgements

We thank all current and past members of the Environmental Genomics group for fruitful discussions and overall support. Research in the group of EHS is supported by the Max-Planck Society, the state of Schleswig-Holstein and the DFG priority program SPP1819.

## Competing interests

The authors declare no competing interests.

## Supplementary Tables and Figures

**S1 Table.** Plasmids, strains and primer designed for this study. Listed are all primers used to create plasmids and probes for Southern blots.

**S2 Table.** Statistics and overview of sequencing data generated in this study.

**S3 Table.** RPKM values of all genes of Zt09 reference and mutant strains. RPKM was calculated using cuffdiff (see Material and Methods).

**S4 Table.** Genes associated to either H3K9me3 or H3K27me3 in Zt09 or mutant strains.

**S5 Table.** Deseq2 results to identify differentially expressed genes between Zt09 reference and mutant strains. Comparisons were performed pair-wise, genes were considered to be significantly different expressed, when │log2 fold-change│ > 2 and padj < 0.001.

**S6 Table.** Enriched GO terms and upregulated genes in the categories DNA replication and RNA-dependent DNA replication.

**S7 Table.** Deseq2 results to analyze expression of transposable elements in Zt09 reference and mutant strains. Comparisons were performed pair-wise.

**S8 Table.** Predicted secondary metabolite gene clusters merged with the *Z. tritici* annotation.

**S9 Table.** Structural variation detected in sequenced progenitor and evolved strains. Listed are location, size and type of structural variation. Only events that have not been described before for Zt09 are listed here.

**S10 Table.** Detailed description and annotation of structural variation detected in the single *kmt1* deletion clones. Some structural variations are associated to large segmental duplications. This is noted as (large segmental duplication).

**S11 Table.** Annotation of breakpoints of segmental duplications in the single clones originating from evolved populations △*kmt1*-50-1 and △*kmt1*-50-2.

**S12 Table.** Distance of structural rearrangements in the evolved single △*kmt1* clones to H3K9me3 (Zt09) and H3K27me3 (Zt09 and △*kmt1*).

## S1 Supplementary Text

Data analysis – programs and commands used for analysis of ChIP-seq, RNA-seq and genome sequencing data.

**S1 Fig. Southern blots to confirm correct integration of deletion and complementation constructs:** for deletion of *kmt1* **(A)**, complementation of *kmt1* **(B)**, deletion of *kmt1* in a *kmt6* deletion background resulting in the generation of a double deletion mutant **(C),** deletion of *kmt6* **(D)**, and complementation of *kmt6* **(E).** Depictured are genomic locations of wildtype (Zt09) and mutant strains, restriction enzymes used, probes, and expected fragment sizes on the blots. All tested strains, except for the <u>underlined</u>, were verified as correct mutants. The strains used for experiments in this study are highlighted in **bold**.

**S2 Fig. Verification of absence of H3K9me3 and H3K27me3 in the respective histone methyltransferase mutant strains.** Shown are the ChIP-seq coverage tracks (normalized to RPKM with deeptools (115) of one replicate per strain. As an example, the coverage of core chromosome 8 **(A)** and accessory chromosome 19 **(B)** is displayed. Based on the missing coverage, we confirm absence of H3K9me3 in the Δ*kmt1* and the Δ*k1/k6* strains and loss of H3K27me3 in the Δ*kmt6* and Δ*k1/k6* strains.

**S3 Fig. Growth assay to compare *in vitro* fitness of mutant strains to Zt09.** All strains were grown in liquid YMS medium at 18°C and the OD_600_ was measured until the stationary phase was reached **(A)**. For each strain, two biological replicates were grown in technical triplicates each. The growth of Δ*kmt1* and Δ*k1/k6* mutants was impaired compared to Zt09 and Δ*kmt6* but was restored in complemented strains. **(B)** We used the R package growthcurver [97] to calculate r-values for each growth curve. The values for Δ*kmt1* and Δ*k1/k6* were significantly lower compared to Zt09, but this was not the case in the complemented strains and Δ*kmt6* (* P ≤ 0.05, ** P ≤ 0.01).

**S4 Fig. Assay to compare tolerance of Zt09 and mutants to different stress conditions** including osmotic stress (NaCl and Sorbitol), oxidative stress (H_2_O_2_), genotoxic stress (MMS, actinomycin D), temperature stress (28°C), cell wall stress (Congo Red), and nutrient starvation (H_2_O agar). We observed almost no differences between Zt09 and Δ*kmt6* strains, whereas Δ*kmt1* and Δ*k1/k6* mutants displayed decreased growth, as observed in the growth rate comparison.

**S5 Fig. Stress assay to compare growth of deletion and respective complementation strains. (A)** The Δ*kmt1* mutants showed overall decreased growth and were particularly sensitive to osmotic stress. These phenotypes were restored in the complemented strains. **(B)** Increased melanization at high temperatures, observed in the Δ*kmt6* mutants, was also reversed in the respective complementation strains.

**S6 Fig. Plant infection phenotypes of Zt09 and mutant strains.** We conducted three independent experiments, including 40 leaves per treatment and using at least two biological replicates per strain. Infection symptoms were evaluated and compared as the percentage of leaf area covered with pycnidia (asexual fruiting bodies) and necrotic lesions within the inoculated leaf areas by manual inspection as well as by automated image analysis of scanned leaves [98]. Symptoms in form of necrotic lesions and pycnidia were quantified after 21 days of infection either manually **(A)** and **(B)** or by automated image analysis of infected leaves **(C)** and **(D)**. Senescence on mock treated leaves was identified as necrosis by the automated image analysis and therefore all treatments, including mock treated leaves, show a high level of necrosis in this analysis. Furthermore, first appearance of symptoms was documented by daily screening of inoculated leaves **(E)** and **(F)**. If no symptoms in form of necrosis or pycnidia appeared during the screening period, no data is shown for those treatments. Virulence of both, Δ*kmt1* and Δ*k1/k6* strains was highly impaired. Δ*kmt6* strains were still able to produce necrosis as well as pycnidia, but the symptoms were reduced compared to Zt09. Wilcoxon-rank sum test was performed to test for significant differences (* P ≤ 0.05, ** P ≤ 0.01, *** P ≤ 0.001).

**S7 Fig. H3K9me3 and H3K27me3 enrichment per chromosome and genes and transposable elements associated with those marks in Zt09 and mutants. (A)** and **(B)** display the percentage of sequence coverage of core and accessory chromosomes with H3K9me3 and H3K27me3 relative to the chromosome length. While there are little differences in the overall coverage with H3K9me3 between Zt09 and Δ*kmt6* (A), H3K27me3 enrichment increases on core chromosomes and decreases on accessory chromosomes in the Δ*kmt1* mutant (B). Chromosome 7 displays a higher H3K27me3 coverage compared to the other core chromosomes as the right arm shows characteristics of an accessory chromosome [31]. **(C and D)** Genes (C) and transposable elements (D) associated with H3K9me3 or H3K27me3 in Zt09 and mutant strains. While there is almost no difference in terms of H3K9me3 associated genes or TEs in the Δ*kmt6* mutants, H3K27me3 moves from genes to TEs in the Δ*kmt1* mutants.

**S8 Fig.** Evolution experiments to monitor genome and chromosome stability during mitotic growth. **(A)** The short-term growth experiment over four weeks assessed stability of accessory chromosomes by screening individual clones in the populations for presence/absence of accessory chromosomes. Strains (Zt09, Δ*kmt1*, Δ*kmt6*, Δ*k1/k6*) were grown in triplicates for four weeks and 4% of the population were transferred to fresh medium every three to four days. After four weeks, single clones were isolated and screened for the presence/absence of accessory chromosomes by PCR. **(B)** A long-term growth experiment over six months was conducted to monitor genome stability in *Z. tritici* populations. Three replicate populations per strain (Zt09, Δ*kmt1*, Δ*kmt6*) were grown in parallel exposed to the same growth conditions. 0.1% of the populations were transferred to fresh medium every three to four days. The genomes of the progenitor strains and all populations after six months of growth were sequenced to detect structural variation.

**S9 Fig.** Pulsed-field gel electrophoresis of mid-size chromosomes of Zt09 and Δ*kmt1* progenitor strains and the three single Δ*kmt1* clones originating from the evolved populations after 50 transfers. While there are no visible differences between the progenitor strains, all three single Δ*kmt1* clones exhibit different karyotypes. Chromosome size marker (M, in Mb) are **Saccharomyces cerevisiae** chromosomes (BioRad, Munich, Germany).

**S10 Fig.** Changes in chromosome structure detected in the evolved Δ*kmt1* clones Δ*kmt1*-50-1-1 and Δ*kmt1*-50-2-1. **(A)** Six duplicated regions were found in Δ*kmt1*-50-1-1, two each on chromosomes 1, 9 and 13. The breakpoints of the first duplicated sequence on chromosome 1 fused to the right telomeres of chromosome 13 (1*13) and 19 (1*19), the second duplication is a tandem duplication. While the right telomere of chromosome 19 fused to chromosome 1, the left arm including the centromere is deleted and *de novo* telomere formation occurred at the breakpoint. Chromosome 13 has two duplicated regions, one is a tandem duplication and one shows *de novo* telomere formation on both ends indicating a breakage of chromosome 13. The right telomere is fused to chromosome 1, the first breakpoint of the first duplicated region provides the new left telomere of the 1*13 chromosome. This breakpoint is located very close to the centromere. The left arm including the centromere forms a new, smaller chromosome 13_1, ending at the right breakpoint of the first duplicated region with *de novo* telomeres. Chromosome 9 did not fuse with another chromosome, but the structural variation rather led to the formation of two smaller chromosomes, both containing the duplicated sequences. The larger chromosome 9 (9_1), ends at the first breakpoint of the first duplicated region with *de novo* telomeres and ends at the breakpoint of the second duplicated region with *de novo* telomeres. The second duplicated region ends at the end of the chromosome where *de novo* telomere formation occurred as a result of chromosome breakage (~12 kb). In the smaller chromosome 9 (9_2), the two duplicated regions fused, deleting the entire sequence between the duplicated regions. **(B)** In the clone Δ*kmt1*-50-2-1, we detected four duplicated regions. While two are located on chromosome 9 and result in a very similar structural variation as described in (A), the other two are found on chromosome 1 and 8. One breakpoint of each duplicated region marks a fusion of the respective chromosomes, while the other one displays *de novo* telomere formation. As a result, three new chromosomes form. A new chromosome that represents a fusion of chromosomes 1 and 8 (1*8) and two chromosomes that are shorter version of chromosomes 1 (1_1) and 8 (8_1). The new, shorter versions both contain the centromeric sequence, while the fused chromosome does not contain any sequences of the original centromeres.

